# Do atmospheric events explain the arrival of an invasive ladybird (*Harmonia axyridis*) in the UK?

**DOI:** 10.1101/681452

**Authors:** Pilvi Siljamo, Kate Ashbrook, Richard F. Comont, Carsten Ambelas Skjøth

**Author notes:** Corresponding author (PS). These authors contributed equally to this work (Pilvi Siljamo, Kate Ashbrook).

## Abstract

Species introduced outside their natural range threaten global biodiversity and despite greater awareness of invasive species risks at ports and airports, control measures in place only concern anthropogenic routes of dispersal. Here, we use the Harlequin ladybird, *Harmonia axyridis*, an invasive species which first arrived in the UK from continental Europe in 2003, to test whether records from 2004 and 2005 were associated with atmospheric events. We used the atmospheric dispersion model SILAM to model the movement of this species from known distributions in continental Europe and tested whether the predicted atmospheric events were associated with the frequency of ladybird records in the UK. We show that the distribution of this species in the early years of its arrival does not provide substantial evidence for a purely anthropogenic introduction and show instead that atmospheric events can better explain this invasion event. Our results suggest that air flows which may assist dispersal over the English Channel are relatively frequent; ranging from once a week from Belgium and the Netherlands to 1-2 times a week from France over our study period. Given the frequency of these events, we demonstrate that atmospheric-assisted dispersal is a viable route for flying species to cross natural barriers.

## Introduction

Species introduced outside their natural range and which have detrimental effects on native species are known as Invasive Alien Species (IAS) and are recognised as a significant component of environmental change worldwide [1,2]. They have been identified as one of the ‘Evil Quartet’ of major drivers of biodiversity loss worldwide [3], are highlighted in the Millennium (2005) [4] and UK National Ecosystem Assessments (2011) [5], and are the focus of Target 5 of the EU 2020 Biodiversity Strategy, including EU regulation 1143/2014 on management of invasive alien species [6]. The direct costs of invasive alien species have been estimated to be approximately US $1.4 trillion, approximately 5% of global GDP [7], with annual costs of £1.7 billion within Britain alone [8].

Many species will arrive in a new country but not establish; of those which do establish most do not become invasive [9–12]. Accurate prediction of the timing, effects and identification of species which may become IAS is not currently possible, despite many attempts [13–22].

Consequently, efforts have focussed on preventing the arrival and establishment of all non-native species. Hulme *et al.* (2008) [2] identified six distinct pathways by which species may spread beyond their native ranges: 1) deliberate release; 2) unintentional escape; 3) unintentional contaminant of another commodity; 4) unintentional stowaway on transport; 5) natural dispersal aided by human-made corridors; and 6) unaided natural dispersal. In line with the 2011-2020 CBD Strategic Plan for Biodiversity, most countries have enhanced their border controls [23] in order to better manage pathways 1-4, but natural dispersal from an introduced population (pathways 5 & 6) is very difficult to police, therefore these pathways are likely to become proportionally more important in the future.

One of the traits associated with successful establishment and invasion, particularly in insects, has been good dispersal ability [24–26], which enables rapid spread beyond the original point of introduction. This is particularly true of the Harlequin ladybird *Harmonia axyridis* (Pallas). Native to eastern Asia, it has been widely introduced outside its native range as a biological control agent and has since spread rapidly to colonise North America, much of Europe, several South American countries, and parts of both northern and southern Africa [27]. The species is a strong flier, actively dispersing over several kilometres to overwintering sites each year [28,29]. It was first recorded in Britain in 2003 and 2004, and it was estimated to have dispersed 105 km per year northwards and 145 kilometres westwards from 2004 to 2008 [30,31].

Once the species was in Britain, it spread northwards and westwards from the arrival point in the south-east, against the prevailing south-westerly wind direction. For this reason, passive wind-borne transport has been considered unlikely to play a major role in its spread [32][30]. However, as the UK is an island, cut off from the rest of the continent of Europe by the North Sea and English Channel, it has been suggested that the original arrival may have been assisted by wind events [32]. This association between atmospheric events and the arrival of this species has not previously been tested.

Here, we investigate whether the arrival of an invasive ladybird, *H. axyridis*, from continental Europe to Great Britain was assisted by atmospheric events, using numerical weather prediction (NWP) models and atmospheric dispersion models (ADM). As a comparison, we also examined human-mediated routes of import: the association of Harlequin records with seaports and airports.

## Material and Methods

### Ladybird records

Biological record data was taken from the UK Ladybird Survey, collated from records submitted to iRecord (www.brc.ac.uk/iRecord), the Harlequin Ladybird survey website (www.harlequin-survey.org) and other data submitted to the scheme. This data is freely available on the National Biodiversity Network at https://registry.nbnatlas.org/public/show/dr695. While, the first three records of the Harlequin ladybird in the UK are from 2003, we focused on records from 2004 – 2005, the main period when the species arrived and established in the UK (139 records in 2004; 2081 records in 2005), Only records submitted with sufficient evidence for independent expert verification (a specimen or adequate photograph) were used in the dataset.

To provide an estimate of the relative proportion of Harlequin records to the background level of ladybird records submitted over time and ensure that spikes in Harlequin numbers were not just good days for recording ladybirds, we compared the Harlequin records to records of six widespread and abundant ladybird species (*Adalia bipunctata* (L.), *Adalia decempunctata* (L.), *Calvia quattuordecimguttata* (L.), *Coccinella septempunctata* (L.), *Halyzia sedecimguttata* (L.) and *Propylea quattuordecimpunctata* (L.)) over the same time period (4479 records in total).

To characterise whether the location of Harlequin records in 2004 and 2005 in the UK (n = 139) were clustered or randomly distributed with respect to ports and airports, we used Ripley’s K-function from package “spatstat” in R. We created two shapefiles, incorporating 59 airports and 31 major ports (Table PORT0103, https://www.gov.uk/government/statistical-data-sets/port01-uk-ports-and-traffic) in England and Wales. To determine whether the Harlequin records were similarly clustered with either airports or ports, we used Monte Carlo simulation with random labelling of points and cross K-function [33].

### SILAM

To simulate atmospheric movements of the Harlequin ladybirds, we used the atmospheric dispersion model SILAM (http://silam.fmi.fi), which is a meso- to global-scale, mathematical-physical atmospheric composition model. SILAM can use both Lagrangian (random walk particle model) [34] and Eulerian (atmospheric dispersion computed in a grid) [35] approaches. It has been used to calculate air quality [36], dispersion of volcanic ash [37], pollen [38] and pest insects [39] as well as dispersion of radioactive nuclides [40] in both forward and inverse (footprint) mode.

The forward atmospheric dispersion models investigate where material will be transported to when the source area is known, whereas the inverse atmospheric dispersion models investigate where the source area is for observed material. Theoretically, forward atmospheric dispersion equations can be used in inverse calculations, only direction of time is negative [34,41]. The forward atmospheric dispersion outputs particle counts, concentration etc. if the source is known well enough or a dispersion area, which describes the area which is affected by the source. The inverse dispersion outputs probability area (∼ footprints), not exact numbers, providing the area where the source could be located within. The source or sources can be located at any point within the probability area and users should evaluate if it is possible (e.g., the source of ladybirds cannot be on the sea). Different data-assimilation methods can reduce the probability area, but they require considerable amounts of observations.

Once the material is emitted to the atmosphere, wind transports particles and gases in the atmosphere, and turbulent eddies mix them. Most of the particles (pollen, dust, flying animals, etc.) which are released inside the atmospheric boundary layer stay there, but some of them are able to escape and go higher. The height of the boundary layer depends on weather, but in mid-latitudes it is typically the lowest 1000-2000 metres of the atmosphere.

Insects generally fly inside the atmospheric boundary layer. This means that atmospheric dispersion models, dedicated to the modelling of particles originating in this layer, are also useful tools to simulate long-distance movements of small insects. As relatively weak fliers, most small insects largely follow the prevailing wind direction and are capable of only slightly affecting their flight directions [42].

The SILAM modelling process requires a source area for the modelled material (particulate matter: PM). Harlequin ladybirds are known to reach a high abundance in urban areas [43], so we took as source areas the larger cities near the north coasts of Belgium (Antwerp, Gent, Bruges, Brussels), the Netherlands (Amsterdam, Hague, Rotterdam) and France (Dunkirk, Lille, Calais, Amiens, Le Havre, Dieppe). Because computational resolution of the model is 0.225° × 0.225° (i.e. about 15 × 25 km), these point sources cover a large proportion of Belgium and the Netherlands. The European Harlequin data was taken from publicly-available data on GBIF (available at https://www.gbif.org/species/4989904). The SILAM forward Harlequin simulations were divided into two parts: sources in France, and sources in Belgium & the Netherlands. This was because the atmospheric wind events would have needed to blow in different directions in order to deposit Harlequins in the relevant sighting areas from the two putative sources.

We modelled dispersal of *H. axyridis* individuals assuming that they would act similarly to coarse particulate matter (particles with a diameter of up to 10 micrometres, known as PM10). Although the ladybird is heavier than this, it has the ability to fly and thus generate its own lift, an advantage not shared by standard particles. We also tested the PM10 dispersal patterns with those for much finer particles up to 2.5 μm in diameter (PM2.5), which generally show very long-range dispersal. This showed very similar dispersal patterns, but PM10 particles dropped out of the airstream more quickly, resulting in concentrations of PM10 lower further from the source areas when compared to PM2.5. Deposition of ladybirds from the airstream was assumed to also be similar to PM10 particles: in particular, rain was assumed to cause deposition.

The first UK records for *H. axyridis* in 2004 were in late June, so we carried out modelling from the 1^st^ June 2004 until 1^st^ October 2004 (the ladybirds largely cease outdoor activity and enter overwintering sites around this time). For 2005, we modelled 1^st^ April-1^st^ October in order to capture both the spring and autumn dispersal periods, as well as the summer activity period. SILAM source points in Europe were modelled as continuously releasing PM10 ladybird particles every day across the two years examined, between 5am and 6pm, UK time.

The ladybird’s habit of overwintering inside buildings causes a spike in records in late autumn as they are noticed by householders. There was no way to reliably split these records from those of ladybirds outside, which might be affected by atmospheric events, so we excluded records from the overwintering period (1^st^ Oct-31^st^ March). We did not model past 2005 as the establishment and rapid spread of the species would have made the distinction between newly-arrived immigrant individuals and existing residents impossibly to quantify.

The ECMWF’s operational numerical weather prediction (NWP) model data (Integrated Forecast System – IFS: https://www.ecmwf.int/) served as a source of weather information both in forward and footprint computations. The NWP model is global and thus it covers the whole dispersion area we were interested in (10.5°W-10°E, 45°N-60°N). We used the data as 3-hour time steps (+3 h, +6 h, +9 h and +12 h forecast lengths) and with a square grid size of 0.225° and 21 NWP-vertical levels from ground to over 5 km. The most important weather parameters used were 3D-winds (the transport of ladybirds) and rain (influences deposition of ladybirds from the atmosphere).

The SILAM harlequin-simulations shown here were computed using Eulerian-SILAM using the same grid as in the NWP model. We used a time step of 15 minutes in the SILAM atmospheric dispersion calculations.

Inverse SILAM (‘footprints’) were computed 72 h backwards from the ladybird observations in 2004. Source points (detection points) and direction of calculation (backward in time) were different than in the forward SILAM simulations, but otherwise the model setup was same. We expected that ladybirds arrived at the earliest one day before observations, latest in the middle of the observation day (e.g., obs. 15/8/2004, “collection time” 14/8/2004 00 UTC-15/8/2004 12 UTC), so the collection time was 1.5 days and simulated period (dispersal time plus collection time) was 1.5-3 days backward. However, the SILAM-footprints showed in many cases that the ladybirds arrived earlier than 1.5 days before observations. Thus, we also computed 10 days inverse simulation for the record on the 30^th^ of June, 2004, which is the earliest UK record in 2004. There the collection time was taken as 10 days as well.

The model output information was produced in 10 km square grid cells. The area extended from the 45°N to the 60°N and from 10.5°W to 10°E to cover the UK, Ireland, Belgium, the Netherlands and part of France, Germany, Denmark and Norway. In case of the forward simulations, the output was given as a daily average, whereas in case of the footprints it was hourly.

### Associating SILAM events with Harlequin arrivals

We used Generalised Linear Models (GLMs) to determine whether SILAM-predicted atmospheric events from source populations were associated with records of *H. axyridis*. If these ladybirds were arriving from cross-channel atmospheric events, we would expect to see *H. axyridis* record numbers to be associated with SILAM-predicted atmospheric events closer to the south-east coastline, with the association reducing with greater distance from the coast. Previous work [44] has found *H. axyridis* able to fly at up to 60km/hr at high altitudes, and to fly for at least two hours. To allow for any extra flight time (Jeffries *et al* stopped monitoring at a two-hour flight time cut-off), plus any short-range flights during the collection period, we split the ladybird dataset into two, and compared *H. axyridis* within 200 km of the continental coastline to records collected further than 200 km from the continental coastline. For both datasets we determined the daily frequency of *H. axyridis* and the daily frequency of the 6 most commonly recorded ladybirds. A bound vector of daily frequency of Harlequins and the daily frequency of common ladybirds was used as the response variable in GLMs with binomial error structures. We calculated the maximum value of SILAM per day from both source populations for a 7-day window for 51°N 1°E (West Kent coastline) around the ladybird record days and used this as an explanatory variable in the models. We were also interested in the association of records with month, and how this differed between years, therefore a combined value of month and year (i.e. 2004.6 to represent June 2004) was also included as an explanatory value.

## Results

### Spatial autocorrelation of airports, ports and Harlequin ladybirds

Bivariate Ripley-K functions suggested that the location of Harlequin records in the first two years of arrival were not associated with airport locations in England and Wales at cluster distances of < 17 km (Fig 1a). Harlequin records were also not associated with port locations in England and Wales from 0 – 5 km cluster distances, but showed some association at greater distances (Fig 1b). As the majority of records of Harlequin ladybirds in 2004 and 2005 are located along the south-east coastline, it is likely that the association at greater distances is due to the existence of several major ports in the south-east (Ramsgate, Dover, Felixstowe, Harwich, Ipswich, and Great Yarmouth).

**Fig 1.**
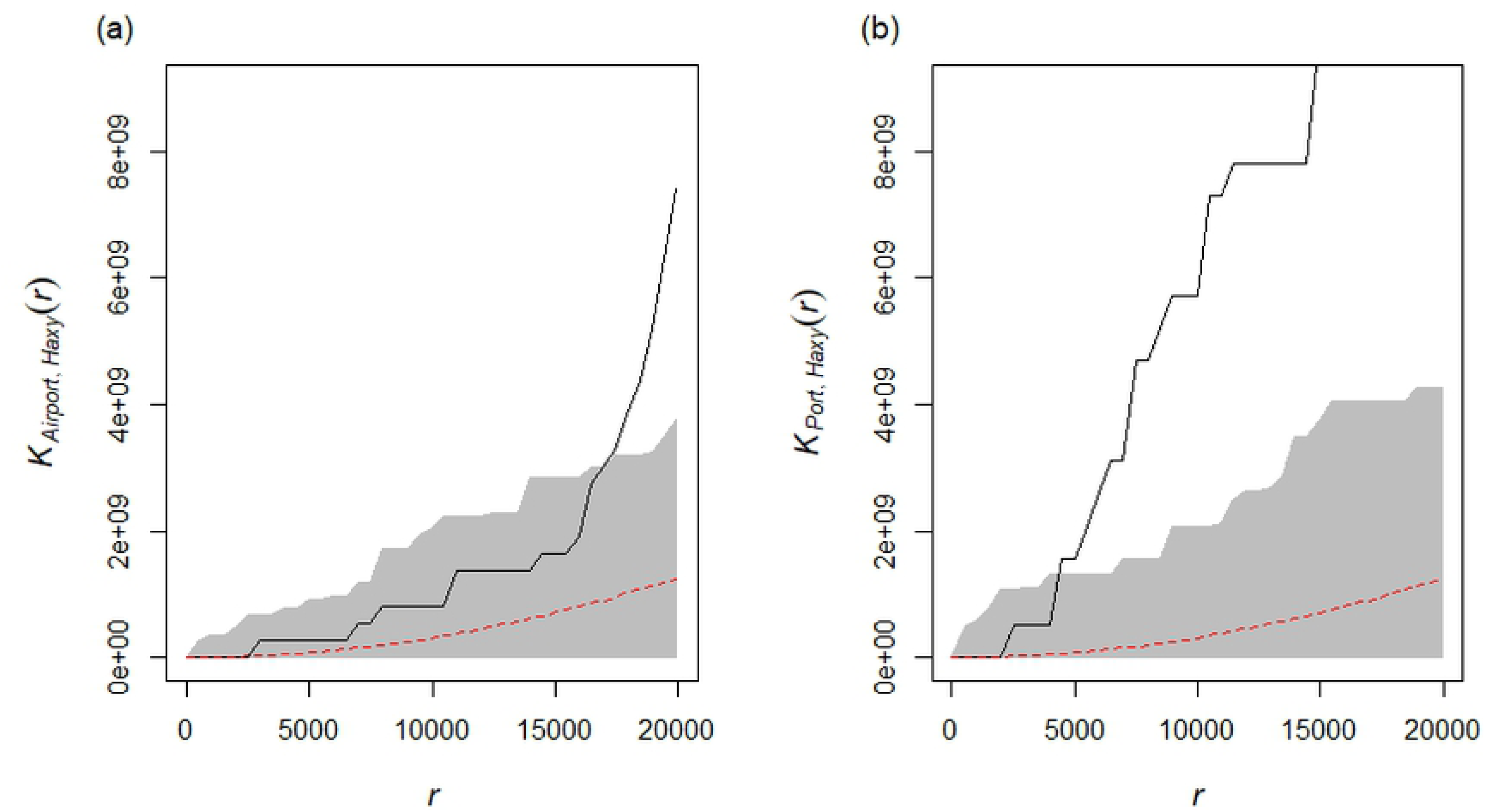
Monte Carlo K-cross simulations for *Harmonia axyridis* records. Monte Carlo K-cross simulations (n=1,000) for *Harmonia axyridis* records in 2004 and 2005 and airports (a) and ports (b). The red dotted line represents what would be expected with the points were randomly distributed; the grey area around this represents the confidence envelope from the Monte Carlo simulations. The black line represents the observed K values; where this line falls within the grey area the points can be described as not associated; however, outside the grey area, the points can be considered to be associated. On the x-axis, r represents cluster distance in metres.

### SILAM events

Air currents from source populations in France, Belgium and the Netherlands could feasibly have transported *H. axyridis* individuals across the English Channel, as the first records are located in the south-east, in line with the SILAM predictions (Fig 2). These predictions suggest that 2005 may have been more favourable for the migration of ladybirds than 2004, as the events were stronger and more frequent in the latter year, especially from Belgium and the Netherlands, where the population appears to be stronger. More favourable winds in the later year can be seen also in Fig 2, where the average potential landing area is larger in 2005 than in 2004.

**Fig 2.**
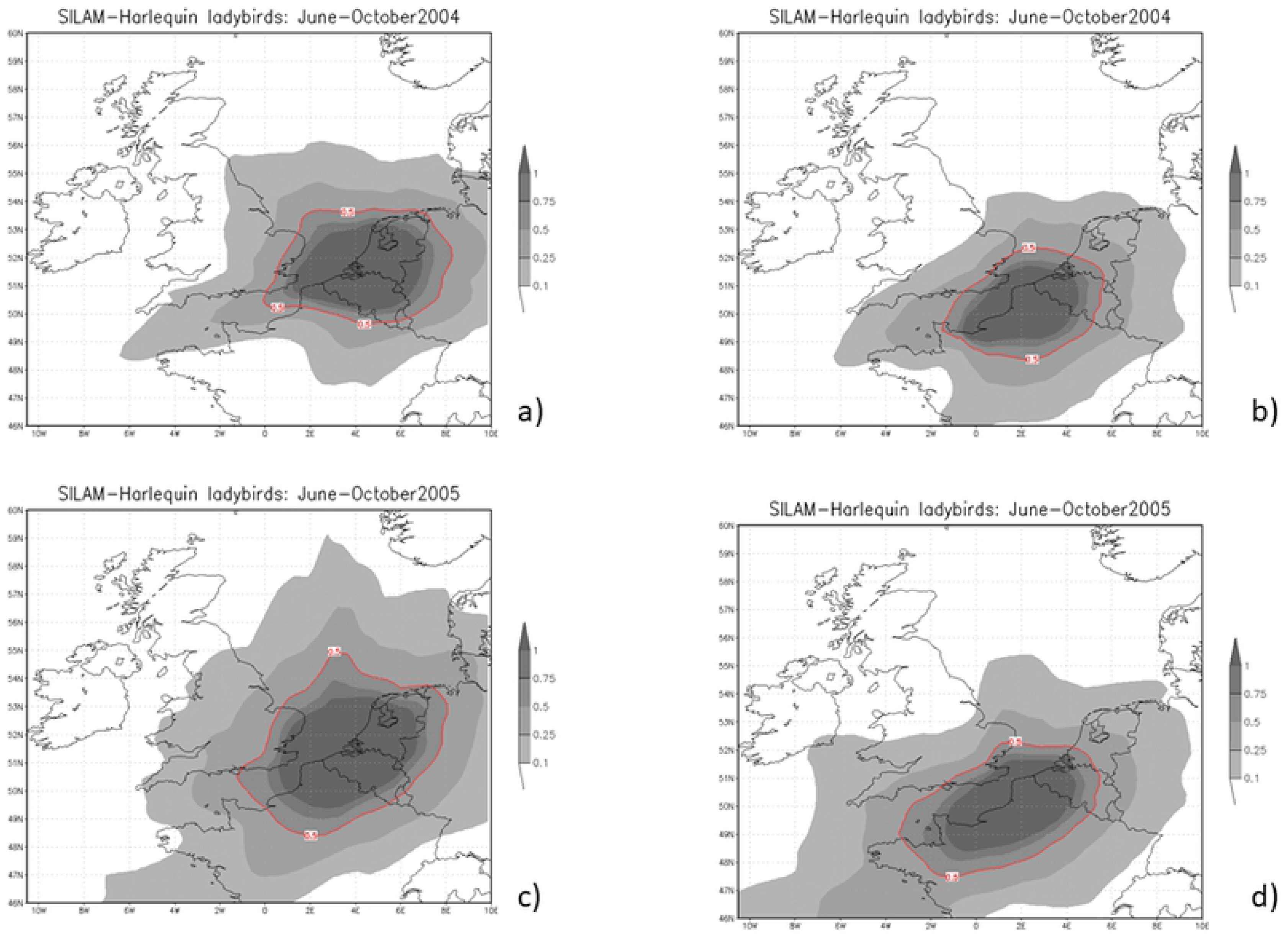
SILAM average air current (PM10) predictions. SILAM average air current (PM10) predictions for June to October 2004 (a,b) and 2005 (c,d) for Netherlands & Belgium combined (a,c) and France (b,d). Higher values (inside contours) suggest a high probability of arrival in the UK from source populations due to atmospheric events.

A more detailed review shows that air flows are suitable for ladybirds to come from Belgium and the Netherlands, on average, once a week (13.9% of days) during June-September 2004, but from France 1-2 times a week (20.5% of days). In 2004, the best month to fly over the English Channel was September (23.3% of the days from Belgium and the Netherlands and 20% from France were suitable). In April-September 2005 a higher proportion of days were suitable for ladybirds to cross the English Channel: 17.6% of the days from Belgium and Netherlands and 22.0% of the days from France. September 2005 was particularly favourable for migration from the continental Europe to the UK, with conditions suitable for immigration from Belgium and the Netherlands almost every four days (23.3%) and more than every three days (33.7%) from France. This coincides with observations of *H. axyridis* becoming more common.

We computed the inverse SILAM (footprints) for all *H. axyridis* records observed before October 2004. The SILAM footprints suggested that France could be the source area in 3 cases out of 7, Belgium or Netherlands in 2 of 7 and the source area is unclear in 2 of 7. In many cases, it is likely that the recorded individual had been present for some time before it was recorded, therefore 1.5 days of collection time only provides a limited snapshot of those records which may be associated with particular atmospheric events, such as the case on 30^th^ June, 2004.

Fig 3 shows an example of the 10 days cumulative, backward probability area from Faversham, Kent (51.3° N, 0.9°E) for 30^th^ June, 2004, which was one of the first *H. axyridis* records in the UK. More detailed analysis shows that the *H. axyridis* observed on 30^th^ June most likely came to the UK during morning hours on 26^th^ June. Several days before and after that short moment winds did not blow from the continental Europe. In this case, the most probable source area locates in France instead of Belgium or the Netherlands.

**Fig 3.**
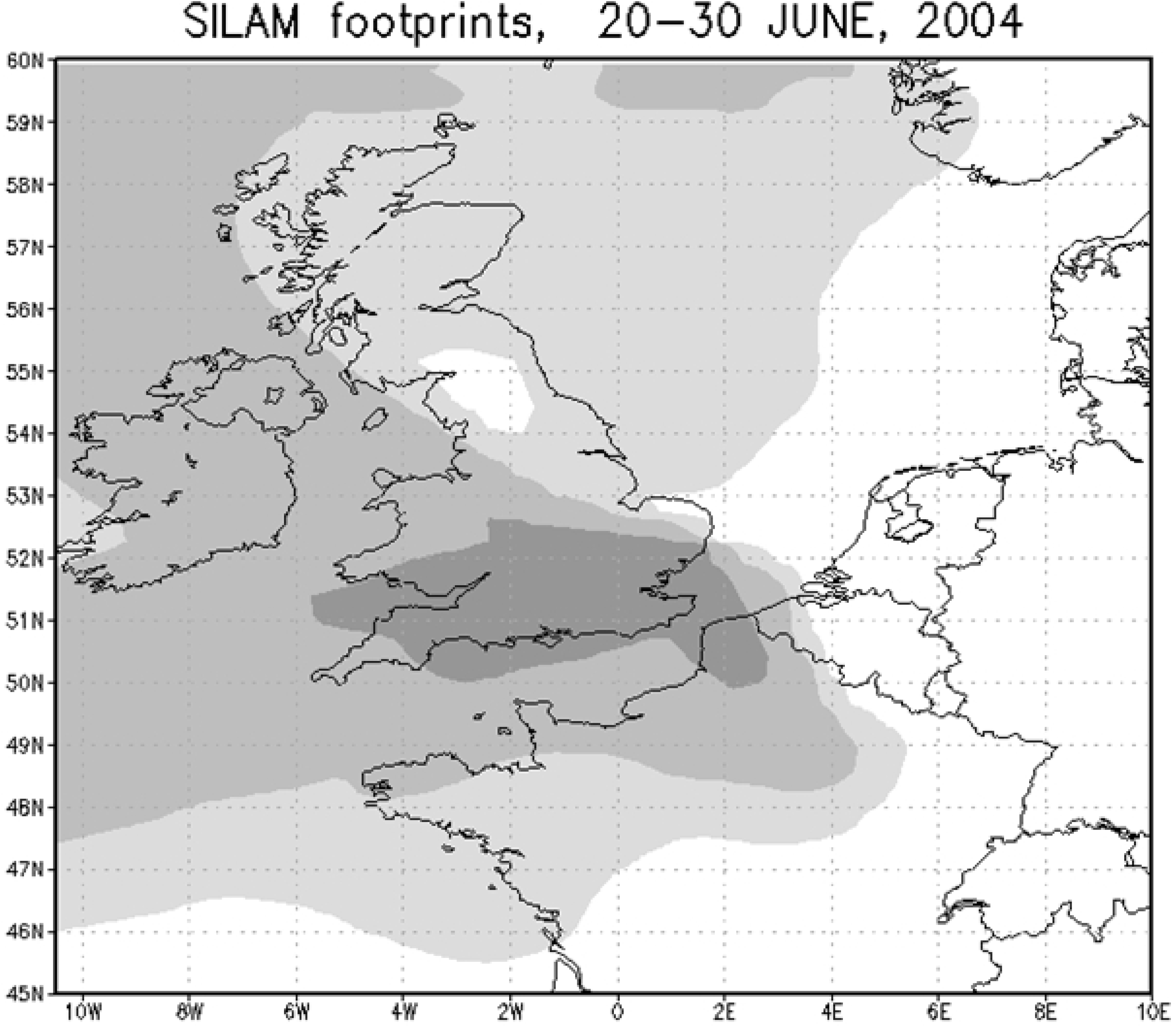
Inverse SILAM simulation. Inverse (footprint) SILAM simulation 10 days backwards from *H. axyridis* record in Kent (51.3° N, 0.9°E) on 30^th^ June, 2004 expecting continuous collection time. The model demonstrates the potential source areas for the record, with darker areas indicating a greater probability of the source location, though it should be noted that ladybirds could only have originated from terrestrial areas.

According to SILAM simulation, *H. axyridis* could fly from France to the UK (Kent and Essex) within 1-3 hours, which is a feasible flying time for ladybirds ([45] p.348). It would take longer (4-6 hours) to reach more northern locations like Suffolk and Norfolk from France.

Another example is for 2005. On 1^st^ Sept, 2005 there were 13 *H. axyridis* records, 72% of all observed ladybirds in the UK that day, all of which were within 200 km of the coastline. The SILAM forward simulations from Belgium and the Netherlands (Fig 4a) and from France (Fig 4b) shows that in this case the source was more likely to be located in Belgium and the Netherlands than in France. The case was also clearly stronger compare to case in the end of June, 2004. Thus, the weather provided an efficient path for Harlequins to arrive in the UK.

**Fig 4.**
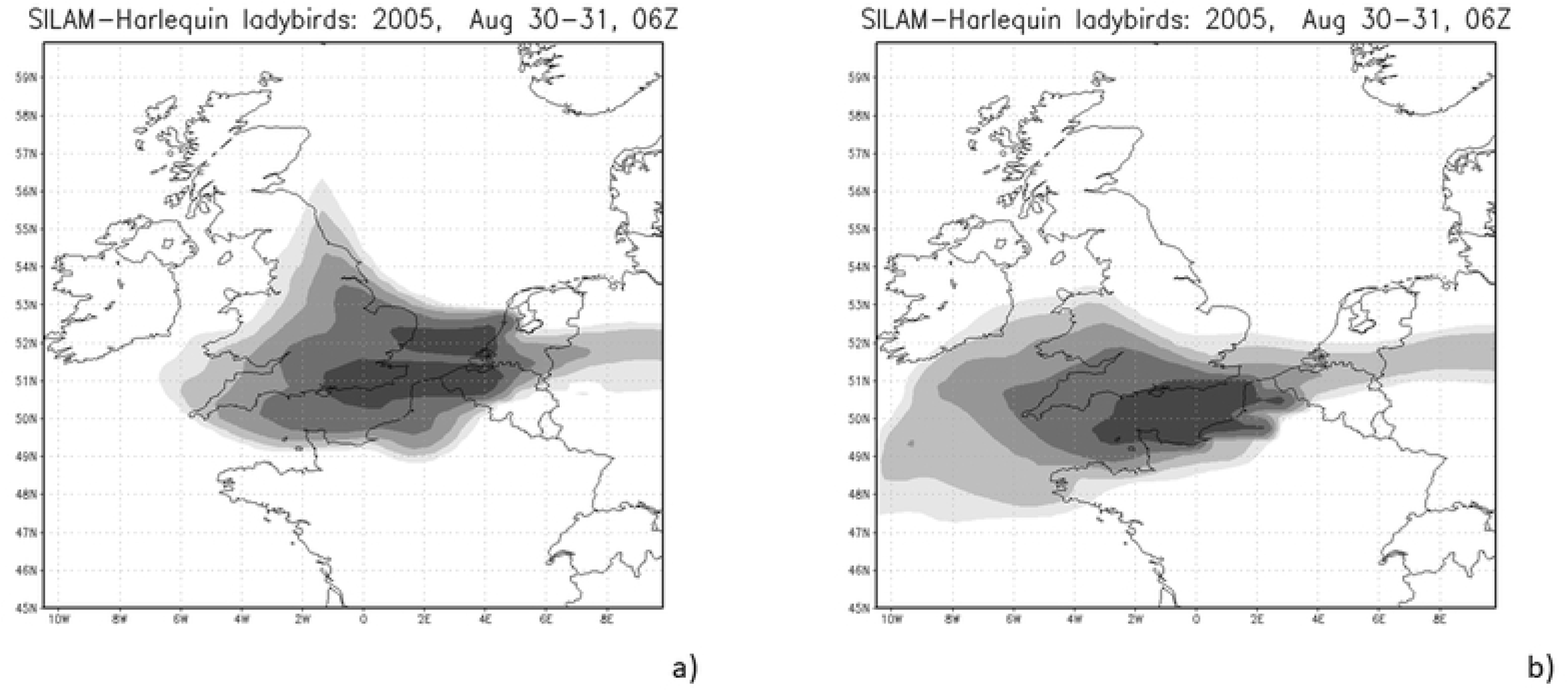
SILAM forward simulations. SILAM forward simulation on 30^th^ August, 2005, 6 UTC - 31^st^ August, 2005, 6 UTC. Source areas locate in a) Belgium and the Netherland and b) in France.

### Association of H. axyridis records and SILAM events

At distances less than 200 km from the continental coastline, we found that there was an association between the daily maximum SILAM values and the proportion of *H. axyridis* records (Quasi-binomial GLM: correlation coefficient: 0.42; LR: 4.22, p = 0.04). With greater values of SILAM, higher proportions of *H. axyridis* records were submitted compared to common native species (Fig 5). There was significant variation in the proportion of *H. axyridis* records to the combined total of the six common native species over time (Fig 5; LR: 31.94, p < 0.001), with a mean proportion of 0.10 *H. axyridis* to native species from June to September 2004, but increasing to a mean proportion of 2.53 *H. axyridis* to native species from April to September 2005.

**Fig 5.**
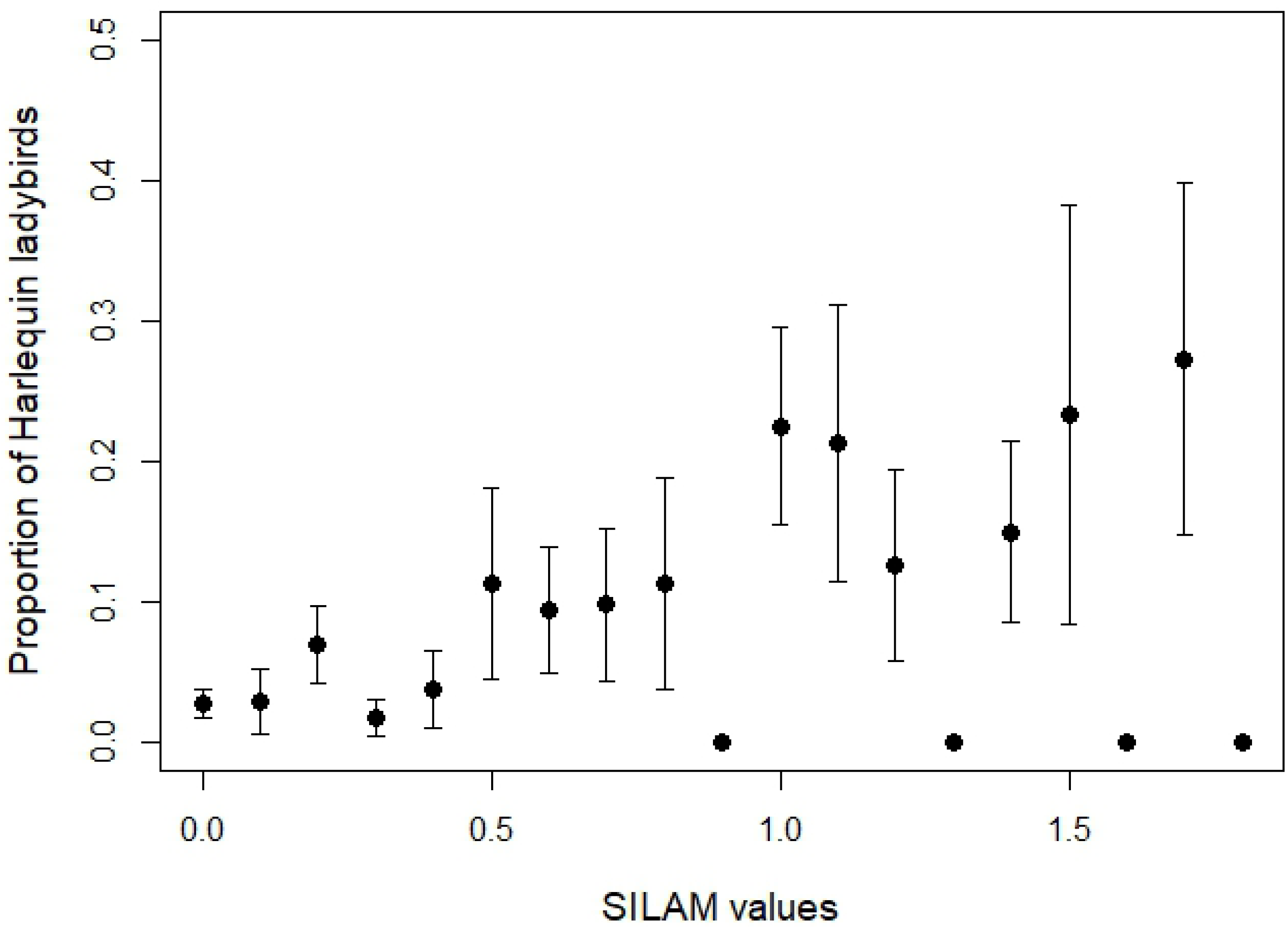
Mean proportion of *H. axyridis* records. Mean proportion of *H. axyridis* records to common species within (black circles) and further than (grey triangles) 200 km from continental coastline with SILAM atmospheric event values. SILAM events have been rounded to nearest 1 decimal place. Error bars represent ±1 S.E.

In contrast, at distances greater than 200 km from the continental coastline, we found that there was no association between the daily maximum SILAM values and the proportion of *H. axyridis* records (Binomial GLM: correlation coefficient: 0.64; LR: 2.79, p = 0.09). However, there was significant variation in the timings of these records (Month & Year: LR: 38.75, p < 0.001), with higher mean proportions of *H. axyridis* to common native species recorded in May 2005 (0.42), June 2005 (0.43) and September 2005 (0.43) than in July 2005 (0.29) and August 2005 (0.26).

## Discussion

Invasive alien species are widely recognised as agents of global biotic homogenisation and thus as one of the main challenges to future global biodiversity [46]. The spread of *H. axyridis* is known to have been assisted by anthropogenic introductions, and some have also alluded to possible atmospheric-assisted dispersal [30,32]. Here, we demonstrate that atmospheric events are a viable, likely and detectable means for this species to have dispersed over a large natural barrier between continental Europe and the UK in 2004 and 2005.

We found some clustering of records of *H. axyridis* with port and airport locations at a regional scale, due to the preponderance of these transport hubs in the south-east near the European continent. However, we found no clustering at small scales, indicating that the species’ records are not clustered in the direct vicinity of ports and airports, as would be the case if these were the primary means of introduction, but were instead scattered more broadly across the southeast of England. It is impossible to rule out the role of anthropogenic transport entirely: indeed, there is considerable anecdotal evidence of ladybirds being moved on ships and other motorised transport [47]. However, the lack of small-scale clustering, combined with the correlation in timing between atmospheric events and ladybird records, suggests that atmospheric transport is a more likely primary method for the species’ arrival.

*Harmonia axyridis* has been reported in 53 countries outside its native range, and when examining the spread of the species it is striking that island nations, and those that have strict detection and prevention systems (e.g. Australia, Cyprus, Iceland, and Malta) unaffected [27,48]. As controls become stricter on anthropogenic transport pathways, it is likely that natural cross-border dispersal from an invasive population will become an increasingly important means of spread for non-native invasive species. Our results indicate that it is possible for *H. axyridis* to be carried across the English Channel and successfully establish: other small winged animals are likely to be able to undertake similarly-assisted dispersal to the UK and other island nations. Indeed, over the last two decades, there have been many new species that have established in the UK: some good fliers, such as the 20 new species of moth [49], but others less associated with flight including the ladybirds *Henosepilachna argus, Rhyzobius chrysomeloides, Rhyzobius lopanthae, Rhyzobius forestieri,* and *Scymnus interruptus* [50].

Wind-assisted passage from continental Europe may be a particularly important route for species that are more adapted to passively utilising wind currents to disperse, such as moths and juvenile spiders. The ballooning behaviour of Wasp spiders *Argiope bruennichi* [51], where individuals are carried by wind-blown silk threads, combined with favourable atmospheric events during the species’ dispersal period, may well have been instrumental in the arrival of this species from continental Europe to the UK in the 1990s. Range-expanding species may also be better-able to exploit favourable weather conditions for dispersal, answering the question of why invasive species newly-arrived in an area can quickly move on elsewhere, but long-time residents do not spread. Higher levels of dispersal by individuals at the edge of populations showing expanding ranges has been demonstrated for *H. axyridis* in Western Europe: Lombaert *et al.* [52] show that there is a rapid increase in individual flight speed over eight years of range expansion. Similar processes have been predicted by several studies of other taxa, e.g., [53–55]. We suggest that the combination of traits associated with greater dispersal potential, together with frequent atmospheric events facilitating long-distance dispersal, is likely to be particularly important for saltatorial population expansion across waterbodies or other large-scale barriers to spread. Once introduced, other factors, such as climate, may play a more influential role in its spread each year [56,57], although this expansion may also be assisted by smaller-scale atmospheric events.

Our results demonstrate that Harlequin ladybirds colonising the UK via atmospheric events had more opportunities to have originated from France, as southerly winds are more common than easterlies. However, from the available evidence, populations of *H. axyridis* appear to be larger in Belgium than in France and so, despite fewer atmospheric events originating from Belgium, each event has a higher likelihood of bearing ladybirds. This probable influx of individuals from multiple sources has likely contributed to the later successful establishment and spread via intraspecific but interpopulational admixture [58,59].

In many invasion events (but not all), source populations can be identified using genetic methods [60]. However, this approach may be affected by sampling errors [61], and it is not predictive in terms of arrival dates or methods. Atmospheric modelling, however, may provide a more rapid assessment of potential source locations when speed is required. Crucially, this approach also provides information on the timing and method of arrival, allowing a better picture of the conditions which facilitate the arrival of non-native species. This data can then be used to predict arrivals, allowing the warning and priming of survey networks, for instance by circulating photographs of the potential arrival with a request for any records which might arise.

One major limitation of this approach is that biological records used within the model, and also those used in model evaluation, need to be relatively comprehensive. Biological recording schemes have increased in presence and reach in the last decade, particularly with the use of online tools, but for novel species there may be a lag between arrival and sufficient records to build an accurate picture of the introduction event. Although there are few Harlequin ladybird records from 2004, it is possible that this species established earlier: three specimens are known from 2003 [32]. As a volunteer survey with (in 2004) relatively limited participation, the network was not particularly sensitive to detecting low numbers of a new species. However, it should be noted that *H. axyridis* is a large and obvious species which often lives in close proximity to people and is apparent even to non-entomologists.

While many countries have tightened their airport security in response to increased knowledge of IAS [23], finding individuals of small species is still challenging, particularly if in personal luggage or live plants. Moreover, individuals assisted by unpredictable atmospheric events to cross large natural barriers bypass these security measures; therefore, a more integrated approach to IAS management should include tracking storm events and subsequent records. These should be used to develop predictive models of periods of high risk of arrival, and surveillance (including working with volunteer recorders) increased. The Harlequin ladybird has had a dramatic impact on native ladybirds in the UK, eating the larvae of many species [27,62,63]; if this invasion had been detected and managed appropriately in the early stages, this may not have occurred.

We hope that the ongoing growth in biological recording, with increasing availability of resources and speed of communication of sightings, will make recording schemes a better real-time early warning system for novel arrivals. More recorders, more and faster access to verifiers, better platforms for timely mass publication of sightings (such as social media), along with greater and more accurate public awareness of novel, potentially harmful species, such as the Asian hornet *Vespa velutina* or Asian Longhorn beetle *Anoplophora glabripennis*, all make speedy detection, identification and dissemination of new species both more possible and more likely. The GB Non-native Secretariat has a list of many potential invaders [61]; if these species are targeted for public awareness campaigns, they may be detected before they establish. One such project currently holding off a full-scale invasion is that concerned with the Asian hornet. In 2004, the Asian hornet *Vespa velutina* was accidentally introduced to south-west France and has spread rapidly, with sightings in Spain [64], Portugal [65], Belgium [66] and Italy [67]). A predator of European honeybees *Apis mellifera*, arrival of this species has been associated with economic impacts on apiculture and pollinator decline [68]. It was first recorded in the UK in 2016 [69,70] and has been found across the south of England from Cornwall to Kent [70]; using storm events to predict areas in need of enhanced nest surveillance may help to reduce the likelihood of this species becoming established in the UK.

Atmosphere is a viable route for invasion, over which we have no control. Given current uncertainty about future climate change, greater frequency of storm events for example, could increase or decrease risk of invasion via this pathway.

## Acknowledgements

The SAPID project of the Academy of Finland supported this study. We would like to thank all the volunteer recorders who have submitted records that made this paper possible, and both David and Helen Roy of the Biological Records Centre who allowed us to use the ladybird dataset. We would also like to thank the many verifiers for the UK Ladybird Survey and the technical support teams at BRC.

